# Regulation of distinct STAT3 dynamics in pro (IFNγ) and anti (IL-10) inflammatory pathways and in their cross-talk: insights from a data-driven model

**DOI:** 10.1101/425868

**Authors:** U Sarma, M Maiti, A Nair, S Bhadange, A Srivastava, B Saha, D Mukherjee

**Affiliations:** National Centre for Cell Science, NCCS Complex, Ganeshkhind, SP Pune University Campus, Pune 411007, India

**Keywords:** IFNγ, IL10, STAT1, STAT3, SOCS1, mathematical modeling, entropy, negative feedback

## Abstract

IFNγ and IL10 pathways elicit functionally opposing cell fates, however, these two stimuli activate common transcription factors like STAT1 and STAT3 (S/1/3). How the two stimuli regulate the dynamics of S/1/3 activation remains less understood. Here, we experimentally measured the signaling dynamics of S/1/3 in response to IFNγ and IL10 and found that STAT3, in particular, exhibits a bell-shaped response to both stimuli with maximal activation in an intermediate dose. We built a mathematical model, which quantitatively captured the S/1/3 dynamics and by model analysis, we identified the primary regulators controlling the bell-shaped STAT3 responses in both pathways. As the STATs are activated in response to both stimuli, in a scenario when cells are subjected to co-stimulation of IFNγ and IL10, an interpathway competition to activate the common substrate is plausible. Additionally, a strong transcriptional induction of SOCS1 (a negative regulator of IFNγ pathway) was observed upon IL10 stimulation, suggesting that IL10 pathway can potentially inhibit the IFNγ pathway via SOCS1 induction. To quantitatively understand the S/1/3 responses to co-stimulation we next simulated the same which predicted STAT3 activation and SOCS1 induction dynamics would robustly remain IL10 driven. Subsequent experiments validated the model predictions. Further, the model analysis identified the primary regulators controlling the robustness of IL10-STAT3 axis in co-stimulation. Together, our data-driven model quantitatively captured the dynamics of the STATs in response to both pro (IFNγ) and anti (IL10)-inflammatory stimuli and identifies distinct regulatory elements controlling STAT3 signaling in individual and co-stimulation conditions.

## Introduction

Information encoded in the dynamics of signaling pathways is frequently observed to shape the course of a spectrum of cellular processes like cell growth, proliferation, apoptosis and developmental lineage commitments [1–12]. For instance, a sustained versus transient dynamics ERK can lead to cell differentiation versus proliferation [8]. In TGFß signaling, long duration SMAD2 activation promotes growth inhibition [13, 14], whereas shorter span of SMAD2 activation is connected to progression of epithelial-to-mesenchymal transition [15]. Studies have shown a direct correlation between the signaling dynamics and gene expression. For example, Lane et al, showed that cell-specific distinct NF-ƙβ dynamics is decoded at the level of gene expression in individual cells [16]. Oncogenic B-Raf mutations and B-Raf inhibitors disrupt the dynamics of ERK signal, which alters cell proliferation and perhaps serves as a precursor to cancer [17], suggesting a connection between signaling dynamics and pathological conditions. Dynamic encoding[18] is also observed in cytokine signaling where the measure of dynamic range was observed as a better determinant of cellular responses, when compared to measured signal strength (basal or hyperstimulated). The dynamics of key signaling components like STATs in response to a single or a combination of stimuli, and the regulators of these signaling events primarily remains unexplored.

Of the two functionally opposing cytokine signaling pathways, interleukin 10 (IL10) activated sustained STAT3 dynamics triggers anti-inflammatory (AIF) responses whereas transient STAT1 activated by interferon gamma (IFNγ) elicits pro-inflammatory (PIF) cellular responses [19, 20]. STAT1 and STAT3 (S/1/3) are canonical responders of the IFNγ and IL10 pathways cues [21, 22]. However, in addition to activation of the canonical signaling axis, these signaling pathways also cross-activate each other’s canonical STATs as a non-canonical component [23, 24].

Earlier studies investigated IFNγ-STAT1 and IL10-STAT3 pathways using both experimental observations [20, 25] and mathematical modeling [26]. IFNγ-STAT1 pathway frequently employs feedback regulator(s) such as SOCS1 and tyrosine phosphatases like SHP-2 to negatively regulate the amplitude and dynamics of active STAT1 [25–29]. In the IL10-STAT3 pathway signal termination occurs via ubiquitination, endocytosis, and degradation of the IL10 receptor 1(IL10R1) [30]. Transcriptionally-induced feedback inhibitors of IL10-STAT3 pathway are not well documented. How such pathways with distinct signal regulatory mechanisms encode incoming information in the dynamics of shared signaling intermediates, remains to be systematically explored. Generally, investigation of dose-dependent dynamics of signaling pathways can lead to identification of non-intuitive emergent features of signaling pathways [31] and thus serve as excellent training and testing datasets for calibrate quantitative mathematical models [32, 33]. Such models can facilitate better understanding of the regulatory mechanisms that underlie the experimental observed behavior [32–35].

In this study, our experiments on Balb/c derived peritoneal macrophages revealed distinct dynamics of S/1/3 in response to different doses of IFNγ or IL10 stimulus. We built a simplified model comprising both IFNγ and IL10 pathways, which quantitatively captured the observed S/1/3 dynamics in response to both stimuli. The model suggested signal inhibition at the receptor level as a common mechanism controlling the dynamics of S/ 1/3 in both pathways. Notably, STAT3 amplitude and dynamic range (measured as area under curve; AUC) was highest for a medium (M) dose of IFNγ or IL10, but, at low (L) and high (H) doses STAT3 responses were inhibited. Through Monte-Carlo sampling and subsequent calculation of the entropy and information content of model variables, we identified the parameters critically determining the bell-shaped STAT3 responses in both pathways. As both pathways activate common STATs and 1L10 stimulation induces SOCS1 expression, co-stimulation (IFNγ + IL10 simultaneously) can result in an inter-pathway competition. We next predicted the effect of co-stimulation on the S/1/3 dynamics which suggested STAT3 and SOCS1 induction dynamics robustly remains IL10 driven during co-stimulation. This prediction was further validated by experimentation. By analyzing the parameter space through Monte-Carlo sampling and by calculating the information content of model variables, we could underpin the parameters controlling the robustness of IL10 driven STAT3 signaling against perturbations such as activation via IFNγ stimuli. Together, through data-driven quantitative modeling, we studied dynamic encoding of functionally opposing information in S/1/3 activation, quantitatively predicted outcome of costimulation experiment and identified distinct sets of regulators controlling two novel aspects of STAT3 signaling: 1. bell-shaped activation in response to both IFNγ and IL10 stimuli, and 2. robustness IL10 driven STAT3 dynamics in co-stimulation.

## Methods

### Experimental protocol

Balb/c derived macrophages were treated with increasing doses of IL10 and IFNγ recombinant ligand. The cells were then lysed and processed for immunoblotting. Dose response studies were the basis of selection of the high (20ng/ml), medium (5ng/ml) and low (1ng/ml) doses of both cytokines, and kinetic studies were performed at these three selected doses.

### Immunoblotting

Cells were treated with the reagents (as indicated). After stimulation the cells were washed twice with chilled PBS, and lysed in NP-40 cell lysis buffer [20 mM Tris (pH 7.5), 150 mM NaCl, 10% glycerol, 1mM EDTA, 1mM EGTA, 1% Nonidet P-40, protease inhibitor mixture (Roche Applied Science, Mannheim, Germany) and phosphatase inhibitor mixture (Pierce)]. The lysates were centrifuged (10,500 rpm, 10 mins) and supernatants were collected. Quantitation of protein was performed using the Bradford reagent (Pierce) and an equal concentration of protein in laemmli sample buffer was loaded on SDS–PAGE. The resolved proteins were transferred onto PVDF (Millipore) membrane. The membranes were blocked using 5% non-fat dried milk in TBST [25 mM Tris (pH 7.6), 137 mM NaCl, and 0.2% Tween 20]. Membranes were incubated with primary antibody overnight at 4°C, followed by washing with TBST. This is followed by incubation of membranes with HRP-conjugated secondary antibody. Immuno-reactive bands were visualized with the luminol reagent (Santa Cruz Biotechnology). The STAT1 antibody we used detected both splice variants of -STAT1 (Tyr701), p91 STAT1α and p84 STAT1β, here we detected STAT1α. The STAT3 antibody we used is bound to tyrosine phosphorylated STAT3 molecules of both isoforms STAT3α(86kDa) and STAT3β (79kDa).

### Reagents

Antibodies specific for p-STATI (Tyr-701), STAT1, p-STAT3 (Tyr-705) and STAT3 were purchased from Cell Signaling Technology (Danvers, MA) and those for SOCS1, SOCS3 and β-actin were from Santa Cruz Biotechnology (Santa Cruz, CA). Soluble mouse recombinant IL10 and IFNγ were procured from BD Biosciences (San Diego, CA). RPMI 1640 medium, penicillin-streptomycin and fetal calf serum were purchased from Gibco®-ThermoFisher Scientific ((Life Technologies BRL, Grand Island, NY). All other chemicals were of analytical grade.

### Animals and cell culture

BALB/c mice originally obtained from Jackson Laboratories (Bar Harbor, ME) were bred in the experimental animal facility of National Centre for Cell Science. All animal usage protocols were approved by the Institutional Animal Care and Use Committee. Studies were performed using 6-8 weeks old mice. 3% thioglycollate-elicited peritoneal macrophages were isolated from Balb/c mice and cultured in RPMI supplemented with 10% fetal bovine serum (FBS). After the cells adhered, they were washed with PBS to remove non-adherent population and maintained for 48 hrs in a humidified CO_2_ incubator at 37°C. The cells were serum starved (by addition of media with 0.2% FBS) for 4 hrs before stimulation.

### Mathematical modeling and analysis of IFNγ and IL10 pathway

Both FNγ and IL10 pathways were built as one model where we preferentially switched on either the IFNγ or IL10 pathways (individual ligand stimulation, depicted in Fig, 2a and 2c) or activated both pathways simultaneously (co-stimulation). Details of the model development, calibration, Monte-Carlo sampling, entropy calculation and model validation steps are elaborated in Supplementary file 1.

## Results

### STAT1 and STAT3 phosphorylation in response to different doses of IL10 and IFNγ

Peritoneal macrophages obtained from BALBc mice were stimulated with increasing doses (0.5, 1.0, 2.5, 5.0, 10.0, 20.0 and 40.0 ng/ml) of either IL10 or IFNγ ligands. We observed: STAT1 phosphorylation increased proportional to the increasing dose of IFNγ stimulation till the 20.0/ml ng dose, followed by a decrease (Fig. 1a), but IL10 induced STAT1 phosphorylation significant increased only at 2.5 and 5 ng/ml of IL10 followed by a decrease (Fig. 1b). STAT3 phosphorylation showed a gradual increase with escalating doses of IL10 (Fig. 1b). STAT3 phosphorylation in response to IFNγ treatment was highest at 10.0/ml ng dose and comparable in the range 2.5-10ng/ml (Fig. 1a). For both IFNγ and IL10 stimulations of 0.5 ng/ml and 1 ng/ml, weak induction of their respective non-canonical STATs (STAT1 for IL10 and STAT3 for IFNγ) was observed; at 5 ng/ml both canonical and non-canonical STATs have high phosphorylation and at 20 ng/ml non-canonical STATs amplitude are inhibited in both the pathway. Based on the dose-response we selected three doses of IFNγ and IL10: low (L), medium (M) and high (H), which are respectively 1.0 ng/ml, 5.0 ng/ml and 20.0 ng/ml, for further investigating the signal dose-dependent dynamics of the STATs in both the functionally opposing pathways.

**Figure 1:**
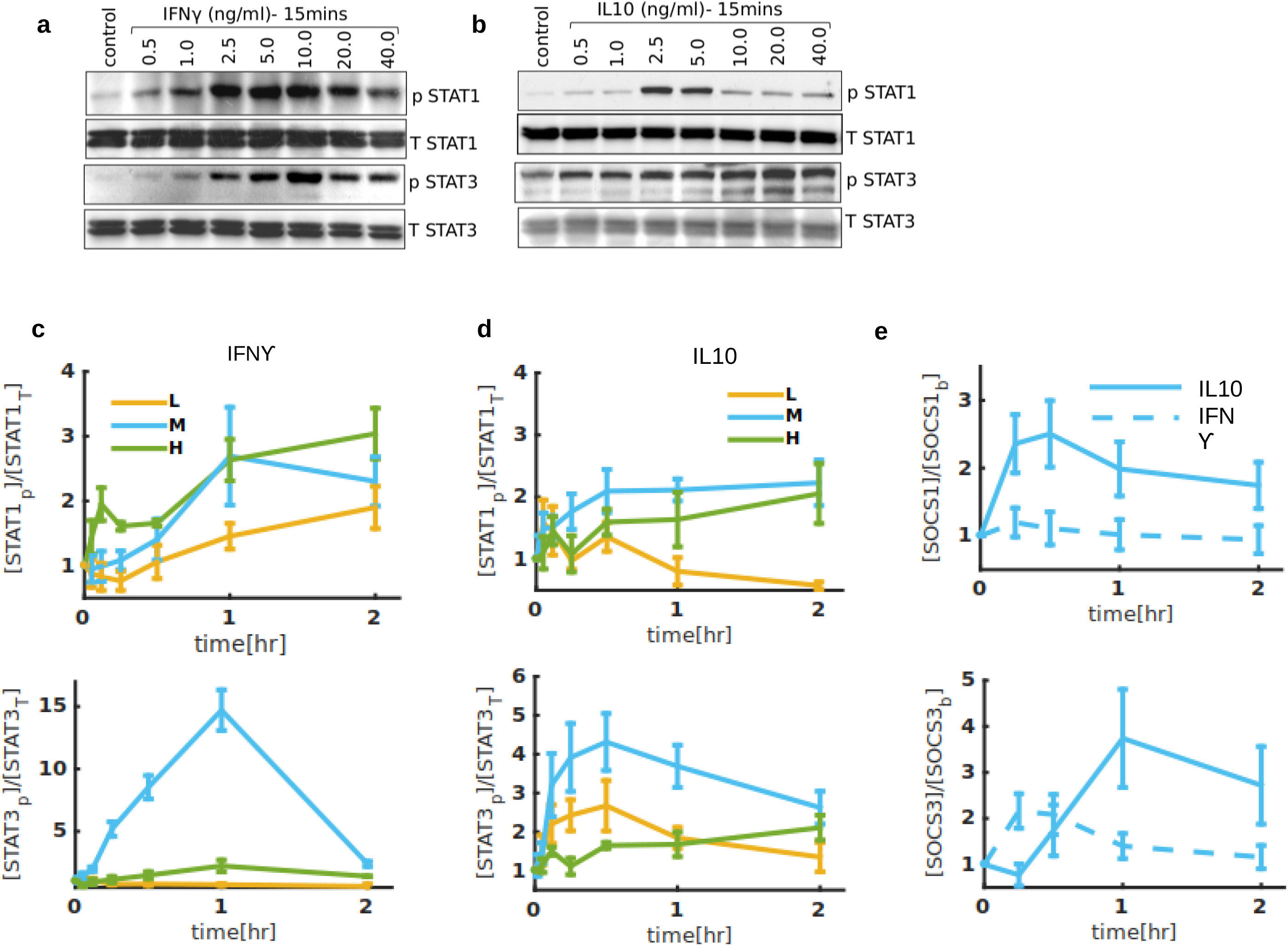
Dose-response and kinetic studies of STAT1 and STAT3 phosphorylation upon IL-10 or IFNγ stimulation. 48 hrs rested Balb/c derived peritoneal macrophages, cultured in RPMI 1640 with 10% fetal bovine serum were subjected to serum starvation for 3hrs. The cells were stimulated with increasing concentrations (0.5ng/ml, 1.0ng/ml, 2.5ng/ml, 5.0ng/ml, 10.0ng/ml, 20.0ng/ml, 40.0ng/ml) of (a) recombinant IL-10 protein or (a) recombinant IFNγ protein for 15mins, then washed with ice-cold phosphate-buffered saline and lysed with RIPA lysis buffer containing protease and phosphatase inhibitors. The cell lysates were further processed for immunoblotting and probed for phosphorylated and total STAT1 and STAT3 proteins. (a) Kinetics of STAT1 and STAT3 phosphorylation by high (20 ng/ml), medium (5 ng/ml) and low (1 ng/ml) doses of IFNγ stimulation.(d) Kinetics of STAT1 and STAT3 phosphorylation by high (20 ng/ml), medium (5 ng/ml) and low (1 ng/ml) doses of IL-10 stimulation. (e) Kinetics of SOCS1 and SOCS3 expression on stimulation with medium dose of IL-10 or IFNγ. Data in (c-e) represents mean ± s.d. of three sets.

Next, cells were stimulated with L, M and H dose of IFNγ and IL10 stimuli and S/1/3 dynamics were captured at different time points, for a duration of 2 hours. For IFNγ treatment STAT1 phosphorylation increased with applied dose strength, although, the change in maximum amplitude and final amplitude were not significantly different (Fig. 1c, 1^st^ panel). However, STAT3 phosphorylation in response to IFNγ stimulation exhibited a distinct dose dependent behavior: at M dose STAT3 is rapidly phosphorylated to a high amplitude that also gets inhibited fast, but at L and H doses STAT3 phosphorylation is suppressed (Fig. 1c, 2^nd^ panel). In response to IL10, STAT1 phosphorylation peaked at M dose with slight inhibition of early activation, but comparable amplitude at two hours was observed at the H dose (Fig. 1d, 1^st^ panel). Notably, a bell-shaped response in STAT3 activation was also observed in response to IL10 (compare Fig. 1d, 2^nd^ panel). Representative immunoblots are shown in supplementary Figure S1. Such dose-dependent inhibition of signal-response implies presence of negative regulator(s) that are shown to inhibit signaling at various stages of the pathways [36, 37]. SOCS1 is a frequently reported negative regulator of IFNγ-STAT1 signaling [26–29], hence we additionally captured dynamics of SOCS1 expression for M dose of IFNγ (Fig. 2e, 1^st^ panel, dashed line). Induction of SOCS1 was also checked in response to IL10 stimulation. Notably, upon IFNγ stimulation, SOCS1 induction remains close to its basal value (Fig. 1e, 1^st^ panel, dashed line), but a stronger SOCS1 induction is observed upon IL10 stimulation (Fig. 1e, 1 ^st^ panel, solid line). Dynamics of SOCS3 induction was also measured as a target gene in both pathways (Fig. 1e, 2^nd^ panel). Studies show, SOCS3 can potentially inhibit IFNγ signaling when SOCS1 is silenced, but the relative inhibitory strength of SOCS3 is negligible compared to SOCS1 when both the SOCS are present in the system [33].

**Figure 2:**
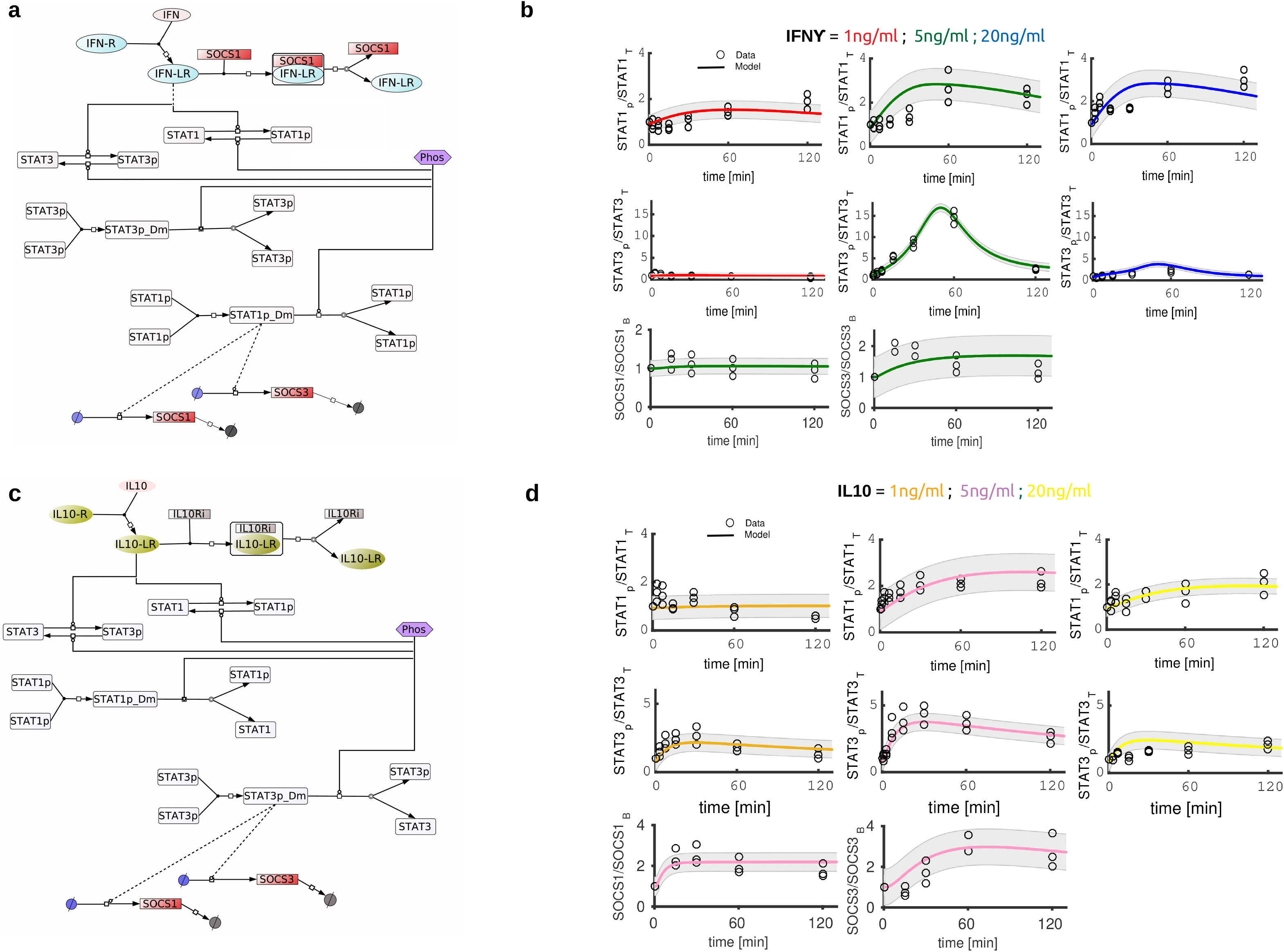
Quantitative modeling of STAT1 and STAT3 dynamics at different doses of IL-10 and IFNγ. (a) Schematic representation of the IFNγ pathway. The blunt heads solid lines representγ pathway. The blunt heads solid lines represent catalysis; arrowheads with solid lines represent binding, unbinding, phosphorylation and dephosphorylation; blunt-headed dashed lines represent transcriptional induction. Upon ligand IFNγ pathway. The blunt heads solid lines represent) binding the receptor IFNγ pathway. The blunt heads solid lines representR) forms an active signaling complex IFNγ pathway. The blunt heads solid lines represent-LR) which triggers the activation of STAT1 (STAT1 → STATp) and STAT3 (STAT3 → STAT3p) through phosphorylation. STAT1p and STAT3p undergo dimerization to become STAT1p_Dm and STAT3p_Dm, respectively. Transcription induction of target genes such as SOCS1 and SOCS3 is shown. SOCS1 is a negative feedback regulator of IFNγ pathway. The blunt heads solid lines representγ signaling which forms a functionally inactive complex [SOCS1.IFNγ pathway. The blunt heads solid lines represent-LR] and inhibits the signaling. Tyrosine phosphatases such as SHP-2 which can dephosphorylate both STAT1 and STAT3 is represented in our model as Phos. (b) IFNγ pathway. The blunt heads solid lines representγ pathway in the model was calibrated to STAT1/3 activation dynamics at three different doses of applied signal 1 ng/ml(L), 5 ng/ml(M) and 20 ng/ ml(H). The pathway was also calibrated to the dynamics of SOCS1 and SOCS3 induction at M. (c) schematics of IL-10 signaling pathways in the model is shown. Nγ pathway. The blunt heads solid lines representotation of the arrows is kept same as described in the IFNγ pathway. The blunt heads solid lines representγ pathway. Both STATs are activated by IL-10 stimulation. Upon ligand (IL-10) binding receptor (IL-10R) forms an active complex (IL-10-LR) which activates both STATs. At the transcriptional level induction of SOCS1 and SOCS3 takes place. IL-10Ri represents an inhibitor of IL-10 signaling which acts by sequestrating and degrading the active IL-10 receptor. (d) IL-10 pathway in the model is calibrated to STAT1/3 phosphorylation dynamics at L, M and H doses of IL-10 stimuli and SOCS1, SOCS3 expression dynamics at M dose. In both (b) and (d) the lines represent model trajectories and blank circles represent respective experimental measurements.

### Quantitative model captures IL10 and IFNγ dose dependent dynamics of STAT3 and STAT1

To understand the regulatory processes controlling the observed S/1/3 dynamics in response to the pro-inflammatory (IFNγ) and anti-inflammatory (IL10) stimuli, we built a mathematical model comprising both the pathways and used the measured dynamics of S/1/3 at different doses of IL10 and IFNγ for model calibration. Both the pathways comprised of three modules

I. Receptor activation module
II. STAT1 and STAT3 phosphorylation module
III. Transcriptional module where SOCS1 and SOCS3 are induced upon IFNγ and IL10 stimulation.

### IFNγ pathway

Fig. 2a schematically shows the structure of the IFNγ pathway as a minimal model. The model has a simplified step of receptor activation that lumps the details of the interaction between IFNγ Receptor (IFN-R) and the Janus kinases JAK1 and JAK2 [26] to a one step binding and activation process [28]. As receptor ligation and complex formation are frequently observed as reversible process [26, 28] we modeled the receptor activation step as a reversible process. Following ligand receptor binding the activated receptor complex (IFN-LR) phosphorylates the transcription factors STAT1 and STAT3. Both the STATs undergo dimerization and forms transcriptionally active complexes [26, 38], which in turn induces target genes such as SOCS1 and SOCS3. Transcriptional induction of target genes was implemented using Hill functions [14]. Dephosphorylation of S/1/3 were assumed to be carried out by constitutively present phosphatases such as SHP2 [25, 39] which is represented as “Phos” in our model.

### IL10 pathway

Fig. 2c shows the schematics of IL10 receptor-mediated phosphorylation of STATs and the transcriptional induction of the SOCS. Similar to the IFNγ pathway, the steps of IL10 receptor 1(IL10R1) and receptor 2 (IL10R2) binding to JAK1, Tyk2 kinases leading to the formation of an active signaling complex [40] were simplified to one step activation-deactivation process. S/1/3 activation was carried out by the active IL10 receptor and their dephosphorylation was assumed to be carried out by constitutive phosphatases such as SHP2 [39]. Negative regulation of IL10 signaling at the receptor level was considered to be carried out by negative regulators like Beta-TrCP-Containing Ubiquitin E3 ligase that binds to and promote degradation of the active IL10 receptor [30].

We studied the responses of STAT1 and STAT3 to L, M and H dose of IFNγ and IL10 by calibrating the model to the measured dynamics of the STATs in each pathway. Details of model calibration steps can be found in Supplementary file 1. The calibrated model quantitatively captured S/1/3 dynamics in response to IFNγ (Fig. 2b, 1st row) and IL10 (Fig. 2d, 1st row) stimulation. The bell-shaped dose response of STAT3 phosphorylation where STAT3 activation is maximum at M dose, but relatively inhibited in both L and H doses of IFNγ (Fig. 2b, 2nd row) and IL10 (Fig. 2d, 2nd row) was successfully captured by the calibrated model. The model also captured SOCS1 and SOCS3 induction dynamics in response to both stimuli (Fig. 2b and Fig. 2d, 3rd row). We measured the SOCS dynamics in M dose as both the canonical (especially STAT3 in IL10 stimulation) and non-canonical STATs are optimally activated in the M doses of both stimuli. Notably, in response to IFNγ stimuli transcriptional induction of SOCS1 was negligible (Fig. 2b, 3rd row), however, basal SOCS1 was required for model calibration wherein SOCS1 negatively regulates IFNγ signaling by blocking the receptor access to its substrate [26]. Notably, SOCS1 expression in response to IL10 was much stronger compared to IFNγ stimulation (Fig. 2d, 3rd row). SOCS3 expression was also captured by the model for both pathway stimulations where SOCS3 was modeled as a target gene in both pathways. During model calibration, the common signaling intermediates and biochemical parameters (between both pathways) were constrained to have a common value such that the shared parameters/concentrations can have one best-fit value that simultaneously captures the STATs and SOCSs trajectories in response to both stimulation. Next, we analyzed the calibrated model to understand the emergence of bell shaped STAT3 responses in both the functionally opposing pathways.

### Regulators of bell shaped STAT3 responses in IFNγ and IL10 pathways

The canonical IFNγ-STAT1 and the IL10-STAT3 pathway exhibited distinct dynamics; STAT1 peak/maximum amplitude in M and H dose are comparable, but STAT3 showed a decline in response at H dose of IL10 and maximum STAT3 activation was observed at M dose. A bell shaped STAT3 activation with much stronger peak was also observed during its non-canonical activation upon IFNγ stimulation. To understand how different variables in the model contribute to the emergence of the observed bell shaped STAT3 responses, and in relation, to identify the key regulator(s), we took the hypercube of best fit model parameters and generated a set of 20000 distinct parameter vectors (in the 0.2-5 fold range of best fit parameters) using Monte Carlo sampling (see details in Supplementary file 1). Simulating the resulting models, each with a distinct parameter vector, we captured the maximum/peak amplitude as well as the area under the S/1/3 trajectories [quantified as area under curve (AUC)], especially focusing on the STAT3 trajectories in response to both IFNγ and IL10 stimulation. The simulations uncover that in addition to the experimentally observed bell-shaped responses, STAT3 also exhibits a different class of signal response in both pathways, where, STAT3 activation increases as a function of signal dose. Fig. 3a and 3c shows the distribution of representative model components such as concentrations of receptors, STATs as well as the induction rate of the negative regulator corresponding to bell-shaped STAT3 responses in both pathways. Similarly, Fig. 3b and 3d shows the distribution of respective parameter sets where bell-shaped response is lost and a proportional STAT3 response is observed. To quantify the parametric contributions underlying a given response type we next calculated the entropy and the information content [41] of each parameter (Supplementary file 1, see information content calculation section). This led to enrichment of the critical parameters whose values are constrained in a specific range to generate a response type such as the bell-shaped STAT3 response. We found, concentrations of IFNγ receptor (IFNR in our model) in the IFNγ pathway and IL10R in the IL10 pathway contains the maximum information regarding bell-shaped STAT3 responses. Moreover, the contribution of other key model variables such as STAT3 concentration itself or the contribution of negative regulators of receptor signaling are relatively much smaller. Further, the maximum information of a single variable (IFNR, ~ 0.8 bits, Fig. 3c) in IFNγ pathway is more than two fold than its counterpart (IL10R) in the IL10 pathway, implying, targeted perturbations of such highly sensitive variable can change the response phenotype more dramatically and flexibly in the IFNγ pathway. The IL10 pathway on the other hand would require simultaneous perturbation of multiple parameters to undergo similar changes. This was subsequently implied in our simulations where on randomly replacing only one/two parameters with maximum information from proportionally responding pathway variants to bell-shaped responders, or vice-versa, we could respectively alter the STAT3 response type with higher frequency in the IFNγ pathway (data not shown).

**Figure 3:**
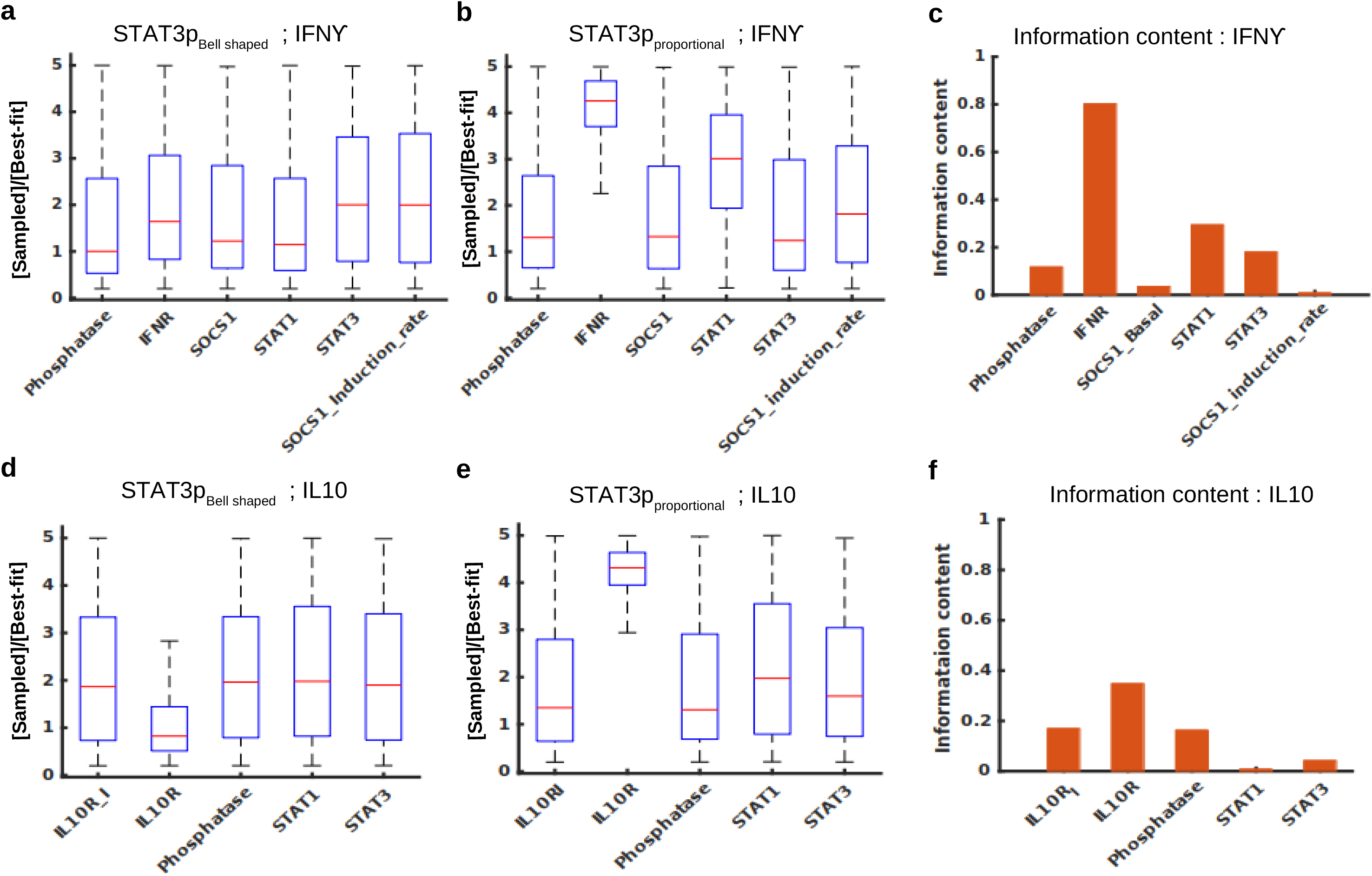
Regulators of bell shaped STAT3 responses in IL-10 and IFNγ pathways. Representative model variables with corresponding to bell-shaped and proportional responder classes are shown as box plots and information content in these parameters is shown as bar plot. Bell shaped (a) and proportional (b) STAT3 responses to IFNγ primarily vary in the expression level of IFNγ receptor (IFNR). Information content analysis (c) quantitatively shows the relative significance of the variables shown in (a)-(b) and amongst all the parameters in the IFNγ pathway IFNR contains the maximum information. In the IL10 pathway the bell-shaped responder classes (d) and the alternate responders generating proportional response (E) also shows sharp differences in IL10 receptor concentration (IL10R) and IL10R contains the highest information (f) in the IL10 pathway variables, however, the absolute information content of IL10R is much less than that of IFNR [compare (c) and (f)].

### Prediction and validation of co-stimulation: STAT3 dynamics remain robustly IL10 driven in presence of IFNγ stimuli

SOCS1 transcriptional induction is negligible in IFNγ pathway although basal SOCS1 acting as a stoichiometric inhibitor of the IFNγ receptor was required for model calibration. However, SOCS1 was found to be strongly induced upon IL10 stimulation (Fig. 2, compare SOCS1 induction in IFNγ and IL10 stimulation). To understand the consequences of IL10 induced SOCS1 on the IFNγ pathway when cells are subjected to co-stimulation we used the calibrated model to predict S/1/3 and SOCS1 induction dynamics in co-stimulation. We chose the M doses of both the ligand types as both the canonical and non-canonical STATs were strongly activated in M doses. The predicted dynamics of STAT3 (Fig. 4a), STAT1 (Fig. 4b) and SOCS1 (Fig. 4c) are shown. As many model parameters are often non-identifiable, to achieve robust predictions we used 40 independently fitted models with comparable goodness of fit [14], which are shown as the shaded area (Fig. 4a-c). The simulations predict, STAT3 signaling would robustly remain IL10 driven when both IL10 and IFNγ stimuli are applied simultaneously. Our experiments subsequent showed that STAT3 dynamics indeed remains strongly IL10 driven (Fig. 4a, experimental data are shown as black filled circles; representative immunoblots are shown in supplementary Figure S2A and S2B). We found a difference in STAT1 amplitudes between model prediction and validation datasets (supplementary Figure S2C), despite the differences in absolute amplitude, shape of the STAT1 trajectory remains closely comparable to the data as a strong quantitative match between model and data was obtained only by adjusting the height of model trajectory with a scaling factor (Fig. 4b, shows the height adjusted model trajectory) while keeping the rest of the model variables unchanged. SOCS1 induction dynamics in co-stimulation also remains comparable to IL10 only stimulation scenario (compare Fig. 4c with Fig. 2d, 3rd row, 1st column). The prediction and validation thus uncovers a feature of IL10-STAT3 signaling: in presence of IL10 stimuli STAT3 dynamics is robust to additional modifications by the IFNγ stimuli, although the latter, when subjected alone, can trigger strong non-canonical activation of STAT3 (Fig. 2b, 2^nd^ row, 2^nd^ column).

**Figure 4.**
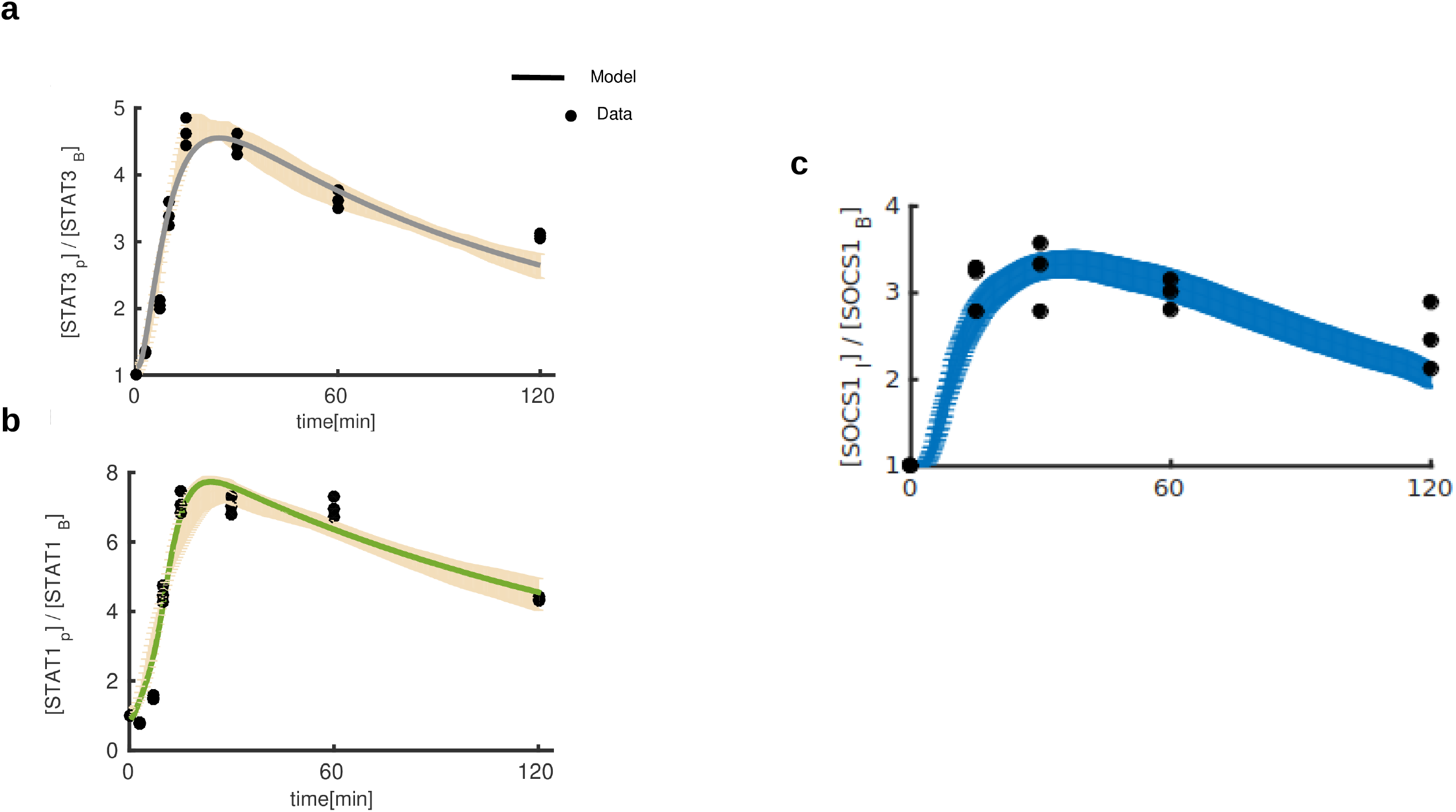
Prediction and validation of STAT1 and STAT3 dynamics in co-stimulation (IL-10 + IFNγ). (a) STAT3 dynamics upon co-stimulation is predicted using multiple best fits with comparable goodness of fit. The shaded area shows predictions from 40 independent best fits and the black dots represent experimental observation.(b) STAT1 dynamics upon co-stimulation is shown, shaded area shows predictions (with amplitude correction using scaling factor) from the 40 independent fits and black dots shows the validation data (c) SOCS1 induction upon co-stimulation. The shaded areas show prediction range and the black dots show experimental measurements. The representative western blots can be found in Figure S2.

### Regulators of robust IL10-STAT3 dynamics in co-stimulation

To underpin the key regulators/variables controlling the dynamics of STAT3 signaling in co-stimulation we sampled the parameter space (Supplementary file 1, Monte-Carlo sampling) and searched for the parameter vectors where STAT3 dynamics is no longer comparable between IL10 and co-stimulation. Specifically, using thousands of sampled parameter vectors we calculated the maximum amplitude and AUC of STAT3 in both IL10 and co-stimulation and indeed our search identified parameter vectors where STAT3 signaling is unique to IL10 and co-stimulation. STAT3 dynamics was considered robust if 0.8 < STAT3_AUC[IL10]_/STAT3_AUC[co-stimulation]_<1.2, and 0.8 < STAT3_max[IL10]_/STAT3_max[co-stimulation]_ <1.2; otherwise STAT3 response was considered as non-robust. Figure 5A and 5B shows the distribution from 5000 parameter vectors each, exhibiting robust and non-robust STAT3 dynamics in co-stimulation. Entropy of individual model variables was calculated (Supplementary file 1) for parameter sets corresponding to both robust and non-robust STAT3 dynamics and information content (Supplementary file 1) of each parameter was extracted. Fig. 5c shows the information content of all the model variables arranged in an ascending order. The parameters in the box plot (Fig. 5a and Fig. 5b) are also shown according to the ascending order of their information content. The analysis uncovered, basal SOCS1 concentration (SOCS1_Basal), binding rate for SOCS1 and activated IFNγ receptor (kf_feedback_IFNg) together with SOCS1 induction by IL10 pathway (SOCS1_induction_rate_IL10) comprises 3 of the top 5 parameters with highest information. Indeed, replacing only these three variables from 5000 robust responders to 5000 non-robust responder systems (with a random selection and replacement from the former to the latter) while keeping the rest of the variables unchanged in the latter, we could obtain more than 50% conversion of the original non-robust responder types to robust responder types (supplementary Figure S3) emphasizing the significance of targeted perturbation of the parameters with high information content.

**Figure 5.**
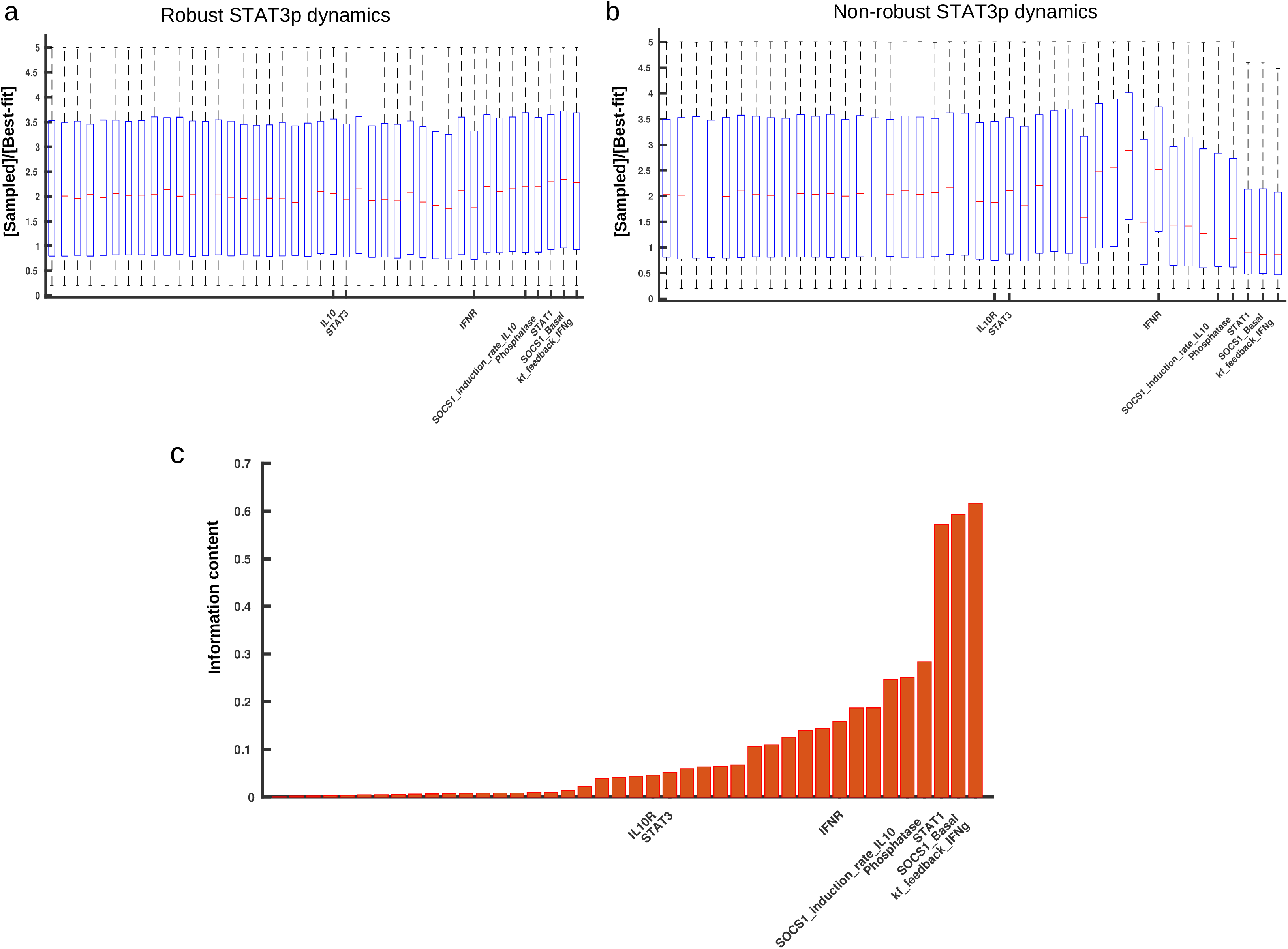
Regulators of robust STAT3 dynamics in co-stimulation. The boxplots show distribution of parameters exhibiting robust (a) and non-robust (b) STAT3 dynamics in costimulation. For each case distributions are shown from 5000 parameter sets sets. Information content was calculated and represented in an ascending order (c). For the sake of comparison, the top 5 parameters with maximum information for STAT3 robustness are labelled together with the parameter that had maximum information for bell-shaped responses (IFNR and IL10R, Fig. 3).

Mechanistically, the reduction/increase in basal SOCS1 concentration has a dramatic effect on the peak amplitude and AUC of the STAT3 trajectory. During co-stimulation the induced SOCS1 inhibits the IFNγ pathway only after the transcriptional time lag, but the early changes in STAT3 activation (as a function of IFNγ pathway basal SOCS1 interaction) resulting from reduced or enhanced SOCS1_Basal dramatically alters the AUC of STAT3 trajectories. The binding rate of SOCS1 and active IFN-LR determines the rate at which IFN-LR is sequestered away as inactive complexes, initially by basal SOCS1, and later by its induced counterpart. Although the IL10 pathway transcriptionally induces SOCS1 the amount of SOCS1_Basal is a key determinant of the observed robustness of IL10-STAT3 signaling axis; induced SOCS1 ensures the maintenance of IL10 driven trajectory further, but, as implied in the information content comparison (Fig. 5), the contribution of IL10 induced SOCS1 in shaping the STAT3 robustness is significantly less compared to its basal counterpart.

## Discussion

STAT1 and STAT3 transcription factors regulate a range of cell fate decisions many of which are functionally opposing. For instance, activation of macrophages is enhanced by STAT1 and inhibited by STAT3, whereas, cell proliferation is inhibited by STAT1 and promoted by STAT3. They act antagonistically in T-helper cell differentiation [42–48]. STAT1 induces the expression of death receptor to promote apoptosis and negatively regulates the expression of several oncogenes [49, 50] while STAT3 activation facilitates survival of many primary tumor cells [49] and majorly activates anti-apoptotic genes that promote proliferation of tumor cells [49, 51]. Cell fate decisions, as observed in multiple studies, are often encoded in the dynamics of signaling proteins. Recent technologies enabled measuring the signaling dynamics and gene expression in the same cells, validating the physiological significance of signaling dynamics in NF-ƙβ pathway [16]. Similarly, dynamics of oncogenic B-Raf [17] or TGFß induced SMAD signaling [14, 15] has distinct cell-fate implications, suggesting that the knowledge of dynamic encoding of signaling information can be critical in understanding the signal decoding process, typically measured in the form of gene expression or different cell fates decisions.

Dynamics of S/1/3 in response to external stimuli is frequently found to represent distinct phenotypic outcomes: IFNγ-STAT1 signaling or IL6-STAT3 signaling is transient and pro-inflammatory whereas IL10-STAT3 signaling is sustained and anti-inflammatory in nature [16, 23]. Both the STATs are however activated in response to IFNγ as well as IL10 stimulation [21–24]. How the functionally opposing information is dynamically encoded in the STATs’ activation remains unexplored. In this study using a combinatorial approach of modeling and experimental data we have addressed the following

1. How the dynamics of canonical and non-canonical STATs are regulated in the IFNγ and IL10 pathway?
2. How functionally opposing cues applied simultaneously would affect the signaling of canonical axis in each pathway?

We conducted experiments in Balb/c derived peritoneal macrophages where we captured the phosphorylation dynamics of STAT1 and STAT3 at different doses of IFNγ and IL10 stimuli. STAT1 responses to different doses of both the stimuli was proportional (or saturating), but intriguingly, STAT3 in response to both stimuli exhibited a bell-shaped response with intermediate (M) doses yielding maximum amplitude and activity (measured as AUC).

To capture the STATs dynamics and to underpin the plausible regulatory mechanisms shaping the observed signal responses we next constructed a simplified mathematical model and calibrated the model to experimental data. The model quantitatively captured the dynamics of S/1/3 in different doses of both stimuli. Next, we used the calibrated model to identify the key variables regulating the bell-shaped STAT3 responses in both pathways. We generated thousands of parameter vectors around the best fit parameter vector and searched for the STAT3 responses which are not bell-shaped. Our analysis identified parameter vectors in both pathways, which no longer exhibited bell-shaped STAT3 responses. Subsequently to determine the key differences between both the observed (bell-shaped) and alternate (proportional) responder types we calculated the information content [41] of the model variables. In both pathways concentration of respective receptors (IFNR in IFNγ pathway and IL10R in IL10 pathway) has maximum information and they critically determine the nature of STAT3 responses. However, the information content of IFNR is two fold higher than IL10R, implying changes in IFNR can more dramatically alter the STAT3 response in IFNγ pathway, whereas, in IL10 pathway similar changes in STAT3 response type would require joint perturbation of multiple model variables with higher information content. We indeed observed this in validatory simulations (data not shown).

Since both stimuli types activates STATs, we next studied a scenario where IFNγ and IL10 compete to activate their canonical signaling axis. This was primarily to understand how the cross-talk between functionally opposing pathways with shared component would regulate signaling through their canonical axis. IL10 stimulation resulted in relatively stronger SOCS1 induction (our experiment shows negligible SOCS1 induction during IFNγ stimulation), indicating additional SOCS1 in the system, if also induced as a function of IL10 stimulation during the co-stimulation, can potentially inhibit IFNγ signaling. We simulated the model for co-stimulation which predicted IL10-STAT3 signaling axis is robust to additional modification by IFNγ stimulus. SOCS1 activation during co-stimulation was also predicted to be IL10 driven. Subsequent experiments quantitatively validated the model predictions. To identify the key variables regulating robustness of STAT3 dynamics we performed Monte-Carlo sampling and generated thousands of parameter vectors that either results in robust IL10-STAT3 signaling or gives rise to distinct STAT3 dynamics for IL10 and co-stimulation. Calculating the information content of parameters we found basal concentration of SOCS1 and binding rate of IFN receptor to SOCS1 (which results in an inactive signaling complex) ensures a robust IL10 driven STAT3 dynamics during co-stimulation, and further, the contribution of IL10 induced SOCS1 is relatively less compared to the basal SOCS1.

Our study explored how two functionally opposing pathways, sharing their signaling intermediates, encoded the input information in the dynamics of signaling. Between the two STATs, our study revealed two novel features of STAT3 signaling during individual (IFNγ or IL10) and co-stimulation conditions. Our quantitative mathematical modeling and subsequent analysis identified key regulators controlling STAT3 signaling during individual and co-stimulation conditions. Future studies may focus on measuring the signaling dynamics (of STAT3 and STAT1) and consequent gene expression profiles in the same cells [16], which can, for instance, directly connect the amplitude and dynamics of IFNγ induced STAT3 to the overall pro-inflammatory gene expression at the single cell level. Additionally, the functional implications of robust IL10-STAT3 signaling may be investigated further to systematically compare if the robustness of IL10 driven STAT3 dynamics (in co-stimulation) is also reflected at the level of genome wide expression of IL10-specific genes.

The intricate balance between and the relative abundance and activation of STAT1 a tumor suppressor and STAT3 an oncogene which are known to have counteracting biological effects, plausibly decides how cells respond to different cytokines as seen in case of IFNγ and IL10 stimulation. In complex and intertwined signaling networks like the STAT signaling, one can take benefit from the systems-level approaches such as the one adopted here and gather a better understanding of the regulatory principles controlling context dependent signaling outcomes. For instance, we could get a better understanding on how two functionally opposing signal cues distinctly shapes the dynamics of a common target such as STAT3. Such cross-talks and resource sharing are frequently observed in the living systems and recent studies indicate plausible evolutionary advantages of such sharing strategies in signaling pathways[41]. Furthermore, systems-level understanding of regulatory mechanisms and their aberrations in pathways critically compromised in diseases such as cancer could open up new avenues to devise better strategies for controlled pharmacological targeting.

## Author Contribution

B.S. and U.S. conceived the study. US built the mathematical model, designed the analysis pipeline and performed the analysis.. D.M. designed and conducted the experiments with M.M, S.G, S.B, A.N and A.S. D.M and U.S interpreted the results. U.S wrote the manuscript with inputs form D.M, A.N. and B.S.

---

**Figure S1. Kinetic studies of STAT1 and STAT3 phosphorylation at low(L), medium (M) and high (H) dose of IL10 and IFNγ stimulation.** Kinetics study of STAT1 (A) and STAT3 (B) phosphorylation at high (20ng/ml), medium (5ng/ml) and low (1ng/ml) doses of recombinant IL10 and IFNγ proteins. Balb/c derived macrophages were stimulated for 3’, 7’, 15’, 30’, 60’ and 120’ with the mentioned doses of IL10 or IFNγ, lysed and processed for immunoblot analysis of STAT1 and STAT3 phosphorylation.(C) Kinetics of SOCS1 AND SOCS3 expression on stimulation with medium dose of IL10 or IFNγ for 15’, 30’, 60’ and 120’ is shown.

**Figure S2. STAT1 and STAT3 dynamics and transcriptional induction of SOCS1 in co-stimulation (IL-10 + IFNγ) treatment.** (A) Representative immunoblots showing measured dynamics of STAT1 and STAT3 activation in co-stimulation is shown for medium dose (5ng/ml) of IL10 and IFNγ. The experimental procedure is same as described in figure S1A or S1B. (B) Kinetics of SOCS1 induction in response to co-stimulation when subjected to medium dose of both IL-10 or IFNγ. (C) STAT1 dynamics predicted from 40 independently fitted models with similar goodness of fit. The shaded area shows the range of predictions and the blue filled circles show the respective experimental data. The model and data trajectories become comparable by multiplying these model trajectory with a scaling factor which results in the trajectory shown in Figure 4C.

**Figure S3. Targeted perturbation of parameters with maximum information changes robustness profile of STAT3 dynamics in co-stimulation.** Figure shows how the parameters with maximum information controls the robustness of STAT3 dynamics. Here, for instance, randomly selecting and replacing only “kf_feedback_IFNg”, the parameter with maximum information, from a set of 5000 robust parameter vectors to a set of 5000 non-robust parameter vectors, while keeping the rest of the parameters constant, we could achieve ~35% conversion of non-robust to robust responses. In the same lines, simultaneously replacing the three parameters related to SOCS1 (kf_feedback_IFNg + SOCS1_Basal + SOCS1_induction_rate_IL10) resulted in more than 50% change of non-robust to robust response types.

**Table TS1:** Table shows the best fit values of model parameters and the upper and lower bound of individual parameters used for fitting. The biological ranges of parameters and species were gathered from literature [26, 28, 38] and the optimization process resulted in the best fit values shown here. Due to the inherent parametric uncertainties involved in experimental measurement of the absolute values of biological parameters we tested the robustness of calibration as well as predictive power of the model using multiple best fit parameters (details in supplementary file 1).

