## Supplementary File 1 for "Regulation of distinct STAT3 dynamics in pro (IFNγ) and anti (IL-10) inflammatory pathways and in their cross-talk: insights from a data-driven model"

Table of content

**A. Building IFNγ and IL10 pathway model**

**AI. Receptor activation and signaling**

**AII. STAT1 and STAT3 signaling**

**AIII. Transcriptional induction of SOCS1 and SOCS3**

**B. Model details**

**BI. Equations**

**BII. Species**

**BIII. Parameters**

**C. Model calibration and validation**

**CI. Datasets for IFNγ pathway calibration**

**CII. Datasets for IL10 pathway calibration**

**CIII. Optimization**

**CIV. Model validation**

**D. Parameter sampling**

**E. Information content calculation**

### A. Building IFNγ and IL10 pathway model

#### AI. Receptor activation and signaling

<u>IFNγ pathway</u>

The ligand (IFNγ) binds to the receptor (IFNR) to form an active signaling complex IFN-LR. We assumed a simple activation-deactivation process in our model[1-3] which simplifies multiple biological steps of receptor signalosome formation .

IFNR + IFN <=> IFN-LR --------- (1)

The signaling complex(IFN-LR) carries out phosphorylation of STAT1 and STAT3(S/1/3). Additionally negative regulators like SOCS1[4] can bind to JAK1[5], a component embedded in the simplified IFN-LR signalosome above and subsequently inhibit the downstream signaling [1, 3, 4] .

STAT1 => STAT1p ; IFN-LR ---------(2)

STAT3 => STAT3p ; IFN-LR ----------(3)

IFN-LR + SOCS1 <=> [IFN-LR. SOCS1] ------ (4)

While in the reactions (2) – (3), IFN-LR acts as a modifier/enzyme, reaction (4) shows inhibition of IFN-LR by the negative regulator SOCS1. The complex [IFN-LR.SOCS1] is functionally inactive which blocks IFNγ mediated phosphorylation of S/1/3 [1].

**B. IL10 pathway**

Similar to the IFNγ receptor the IL10 receptor also binds to its ligand and become functionally active. In our receptor activation module explicit receptor activation deactivation steps are simplified to a one step activation and deactivation process .

IL10R + IL10 <=> IL10-LR --------- (5)

The active signaling complex IL10-LR phosphorylates and activates STAT3 and STAT1. Stoichiometric inhibitor that typically acts by targeting the IL10R1 [5] is modeled as a potential negative regulator of IL10 signaling.

STAT1 => STAT1p ; IL10-LR ---------(6)

STAT3 => STAT3p ; IL10-LR ----------(7)

IL10-LR + IL10Ri <=> [IL10-LR . IL10Ri] ------ (8)

As observed[5], IL10Ri production and degradation is considered as a constitutive process that is independent of IL10 stimulation which are represented as the reactions below

=> IL10Ri --------- (9)

IL10Ri => --------- (10)

Where reaction (9) and (10) depicts the basal production and degradation of IL10Ri.

**II. STAT1 and STAT3 signaling**

STAT1 and STAT3 are activated during both IFNγ and IL10 stimulations.

STAT1 => STAT1_act ; IFN-LR , IL10-LR ----- (11)

STAT3 => STAT3_act ; IFN-LR , IL10-LR ----- (12)

Studies show STAT1 and STAT3 compete with each other for the same receptor phosphotyrosine motif during IFNγ signaling [7] which is implemented in our model as a competition between both the STATs to get phosphorylated by the IFNLR. In the same lines, we implemented competition of STAT1 and STAT3 for access to IL10.

STAT1_act and STAT3_act are the activated/phosphorylated forms of STATs which subsequently undergo dimerization and become transcriptionally active[1]

STAT1_act + STAT1_act -> STAT1_act_Dm ----- (13)

STAT3_act + STAT3_act -> STAT3_act_Dm ----- (14)

#### III. Transcriptional induction of SOCS1 and SOCS3

Transcriptional induction of SOCS1 and SOCS3 were experimentally measured for both IFNγ and IL10 signaling. Both the SOCS are induced relatively strongly upon IL10 signaling compared to IFNγ signaling (Figure 2B and 2D), especially SOCS1 induced upon 1L10 signaling is ~3 fold higher compared to IFNγ signaling.

In our model, in the absence of external signal basal production and degradation of the SOCS is given as

=> SOCS1 ----- (15)

=> SOCS3 ----- (16)

SOCS1 => ----- (17)

SOCS3 => ----- (18)

Where reaction (15)&(16) represents production and reaction (17) & (18) represents degradation of the SOCS.

Studies show upon IFNγ and IL10 signaling both the SOCS are transcriptionally induced as a function of canonical STAT1 (in IFNγ pathway) or STAT3(in IL10 pathway). In STAT1 null mice SOCS3 but not SOCS1 is induced and in STAT3 null mice SOCS3 induction is blocked [7]; thus we modeled SOCS1 and SOCS3 induction as functions STAT1 dimer(STAT1 _act_Dm) and STAT3 dimer (STAT3_act_Dm) respectively. Additionally, studies show efficient promoter binding and gene expression by STAT1 and STAT3 also depends on other complex factors like availability of other cofactors, for instance, during IFNγ stimulation, occupation of DNA-binding sites for STAT1 and the transcriptional activator Sp1 are both required for full activation of certain genes [8]. Similarly gene expression in IL10 pathway are dependent on sp1 and sp3 cofactors [9]. Also competition of bind to different cofactors emerge between STAT1 and STAT3 [10]. Considering the complexity of such interactions and to accommodate the plausible differences in STAT1(STAT3) mediated induction of SOCS1(SOCS3) in IFNγ and IL10 pathways we assumed differences in their induction rates, Km values and hill coefficients and estimated these values during model calibration.

### B. Model details

#### BI. Equations

The differential equations below captures the information propagation in both IFNγ and IL10 pathways. The xs' are model species and the ps are model parameters, names of the species and parameter as well as their best fit values are detailed in supplementary table TS1.

$$\frac{d[x1]}{dt} = x3*p2 - x1*x2*p1 * p0$$

$$\frac{d[x2]}{dt} = x3*p2 - x1*x2*p1$$

$$\frac{d[x3]}{dt} = x1*x2*p1 - x3*p2 + x4*p4 + x4*p16 - x3*x9*p3$$

$$\frac{d[x4]}{dt} = x3*x9*p3 - x4*p16 - x4*p5 - x4*p4$$

$$\frac{d[x5]}{dt} = p8*x6*p7g + p8*x11*p7g - (x3*x5*p6)/(p9*(x5/p9 + x7/p10 + 1)) - (x15*p21*x5*p26)/(p28*(x5/p28 + x7/p29 + 1))$$

$$\frac{d[x6]}{dt} = p8*x11*p7g - p8*x6*p7g - 2*x6^2*p11 + (x3*x5*p6)/(p9*(x5/p9 + x7/p10 + 1)) + (x15*p21*x5*p26)/(p28*(x5/p28 + x7/p29 + 1))$$

$$\frac{d[x7]}{dt} = p8*x8*p14g + p8*x12*p14g - (x3*x7*p13)/(p10*(x5/p9 + x7/p10 + 1)) - (x15*p21*x7*p27)/(p29*(x5/p28 + x7/p29 + 1))$$

$$\frac{d[x8]}{dt} = p8*x12*p14g - p8*x8*p14g - 2*x8^2*p12 + (x3*x7*p13)/(p10*(x5/p9 + x7/p10 + 1)) + (x15*p21*x7*p27)/(p29*(x5/p28 + x7/p29 + 1))$$

$$\frac{d[x9]}{dt} = p15 - x9*p16 + x4*p4 + x4*p5 - x3*x9*p3 + (p19*p15*(x11/p30)^n2)/((x11/p30)^n2 + 1) + (p21*p34*p15*(x11/p32)^n1)/((x11/p32)^n1 + 1)$$

$$\frac{d[x10]}{dt} = p17 - x10*p18 + (p20*p17*(x12/p31)^n22)/((x12/p31)^n22 + 1) + (p21*p35*p17*(x12/p33)^n11)/((x12/p33)^n11 + 1)$$

$$\frac{d[x11]}{dt} = x6^2*p11 - p8*x11*p7$$

$$\frac{d[x12]}{dt} = x8^2*p12 - p8*x12*p14$$

$$\frac{d[x13]}{dt} = x15*p21*p23 - x13*x14*p21*p22$$

$$\frac{d[x14]}{dt} = x15*p21*p23 - x13*x14*p21*p22$$

$$\frac{d[x15]}{dt} = p21*x16*p25 - x15*p21*p23 + p21*x16*p37 - x15*x17*p21*p24 + x13*x14*p21*p22$$

$$\frac{d[x16]}{dt} = x15*x17*p21*p24 - p21*x16*p25 - p21*x16*p37 - p21*x16*p38$$

$$\frac{d[x17]}{dt} = p21*p36 + p21*x16*p38 - x17*p21*p37 + p21*x16*p25 - x15*x17*p21*p24$$

The model above is calibrated to the experimental data (Figure 2), and the calibrated model is used for making predictions that were validated experimentally(Figure 3). Details of model calibration and validation steps can be found in the sections below.

**BII. Species**

| Species | Name in model | Biological meaning |
|---|---|---|
| x1 | IFNγ | IFNγ ligand |
| x2 | IFNR | IFNγ receptor |
| x3 | IFN-LR | Active signaling complex ; IFNγ pathway |
| x4 | IFN-LR.SOCS1 | Functionally inactive signaling complex |
| x5 | STAT1 | Inactive STAT1 |
| x6 | STAT1_act | Active STAT1 |
| x7 | STAT3 | Inactive STAT3 |
| x8 | STAT3_act | Active STAT3 |
| x9 | SOCS1 | Free SOCS1 |

| | | |
|---|---|---|
| x10 | SOCS3 | Free SOCS3 |
| x11 | STAT1_act_Dm | Active STAT1 dimer |
| x12 | STAT3_act_Dm | Active STAT3 dimer |
| x13 | IL-10 | IL-10 ligand |
| x14 | IL-10R | IL-10 receptor |
| x15 | IL10-LR | Active signaling complex; IL-10 pathway |
| x16 | IL10-LR.IL10Ri | Functionally inactive signaling complex |
| x17 | IL10Ri | Inhibitor of IL10R |

**BII. Parameters**

| Parameters | Name in model |
|---|---|
| p0 | IFNγ_On |
| p1 | kf_Active_Rec_IFNγ |
| p2 | kb_Active_Rec_IFNγ |
| p3 | kf_feedback_IFNγ |
| p4 | kb_feedback_IFNγ |
| p5 | kdeg_Active_Rec_IFNγ |
| p6 | kphos_STAT1_IFN |
| p7 | kdephos_STAT1_IFNγ |
| p8 | Phosphatase |
| p9 | Km_STAT1_phos_IFN |
| p10 | Km_STAT3_phos_IFN |
| p11 | kDm_STAT1 |
| p12 | kDm_STAT3 |
| p13 | kphos_STAT3_IFNγ |
| p14 | kdephos_STAT3_IFNγ |
| p15 | kprod_SOCS1 |
| p16 | kdeg_SOCS1 |
| p17 | kprod_SOCS3 |
| p18 | kdeg_SOCS3 |
| p19 | index_k_STAT1_induced_prod_SOCS1 |
| p20 | index_k_STAT1_induced_prod_SOCS3 |
| p21 | IL10_on |
| p22 | kf_Active_Rec_IL10 |
| p23 | kb_Active_Rec_IL10 |
| p24 | kf_feedback_IL10 |
| p25 | kb_feedback_IL10 |
| p26 | kphos_STAT1_IL10 |
| p27 | kphos_STAT3_IL10 |
| p28 | Km_STAT1_phos_IL10 |
| p29 | Km_STAT3_phos_IL10 |
| p30 | Km_STAT1_SOCS1 |

| | |
|---|---|
| p31 | Km_STAT1_SOCS3 |
| p32 | Km_STAT3_SOCS1 |
| p33 | Km_STAT3_SOCS3 |
| p34 | index_k_STAT3_induced_prod_SOCS1 |
| p35 | index_k_STAT3_induced_prod_SOCS3 |
| p36 | kprod_Feedback_inhibitor |
| p37 | kdeg_Feedback_inhibitor |
| p38 | Induced_deg_IL10_receptor |
| p39 | n1 |
| p40 | n11 |
| p41 | n2 |
| p42 | n22 |

**C. Model calibration and validation**

To estimate the variables in the model that best explains the experimental data we fitted the dynamics of S/1/3 trajectories to the corresponding experimental measurements. We have used 16 independent experimental measurements for model calibration and three independent measurements were used to validate model predictions. As both IFNγ and IL10 pathways were built within one single model we selectively activated either of the pathways for a given stimulus condition while fitting the pathways specific experimental conditions. Only during the costimulation predictions both pathways were activated simultaneously. The calibration datasets are listed below

**CI.** Datasets for IFNγ pathway calibration

1. STAT1 phosphorylation kinetics at low dose(L)

2. STAT1 phosphorylation kinetics at medium dose(M)

3. STAT1 phosphorylation kinetics at high dose(H)

4. STAT3 phosphorylation kinetics at L

5. STAT3 phosphorylation kinetics at M

6. STAT3 phosphorylation kinetics at H

7. SOCS1 induction kinetics at M

8. SOCS3 induction kinetics at M

**CII.** Datasets for IL10 pathway calibration

1. STAT1 phosphorylation kinetics at low dose(L)

2. STAT1 phosphorylation kinetics at medium dose(M)

3. STAT1 phosphorylation kinetics at high dose(H)

4. STAT3 phosphorylation kinetics at L

5. STAT3 phosphorylation kinetics at M

6. STAT3 phosphorylation kinetics at H

7. SOCS1 induction kinetics at M

8. SOCS3 induction kinetics at M

### CII. Optimization

Observables: Optimization process involved fitting observables from the model which we fitted against the experimental data. The following observables were subjected to fitting

$$\text{STAT1\_phos} = \frac{(x6+2*x11)}{(x5+x6+2*x11)} \quad \text{--------} \ (1)$$

$$\text{STAT3\_phos} = \frac{(x8+2*x12)}{(x7+x8+2*x12)} \quad \text{--------} \ (2)$$

$$\text{SOCS1}_{\text{IFN}\gamma} = \frac{(x4+x9)}{x9[basal]} \quad \text{--------} \ (3)$$

$$\text{SOCS1}_{\text{IL10}} = \frac{x9}{x9[basal]} \quad \text{--------} \ (4)$$

$$\text{SOCS3} = \frac{x10}{x10[basal]} \quad \text{--------} \ (5)$$

The observables STAT1_phos and STAT3_phos captures the dynamics of phosphorylated STAT1 and STAT3, respectively. Similarly the observables SOCS1 and SOCS3 compares the fold change of induction over the basal The best fit parameters with their upper and lower bound are shown in supplementary table TS1.

Offsets and scaling factors**:** During model calibration we introduced offset and scaling factors which facilitates quantitative comparison of model and data by adjusting the model trajectories without influencing the biologically significant observables and general conclusions offered by the model [11, 12]. To consider the contribution of uncertainties in experimental measurement process and additional unknown smaller technical fluctuations on the day of the experiment we have implemented local scaling factors. In our model the optimization process minimizes the differences between the model and the data from 16 datasets simultaneously to estimate the goodness of fit ($\chi^2$)

$$\chi^2 = \frac{\left(offset_{global} + scale_{global} * scale_{local} * model - data\right)^2}{error^2} \quad \text{----(6)}$$

Optimization was carried out using the MATLAB optimization toolbox Data2Dynamics [13] where goodness of fit (function (6)) is calculated by evaluating a log likelihood function that minimizes the difference between model and data, as well as the contribution from the fitted error models from different data sets. Details of fitting processes can be found elsewhere [13, 14]. Fitting was done using a multi-start local optimization strategy where the starting parameter vectors were generated using Latin hypercube sampling method and MATLAB function lsqnonlin was used for optimization [13, 10]. We estimated one offset_global and one scale_global for all the experimental conditions across the doses or stimulus type applied. For example, one offset_global for STAT1phos is calculated for L,M, H doses of both IFNγ and IL10 treatments. Similarly one scale_global is calculated and in each independent dataset (L, M or H) a scale_localis calculated(in the range 0.7-1.5) to accommodate plausible measurement fluctuations local to individual datasets; this also ensures that overfitting plausibly arising due to scaling factors across all datasets is minimized. The bestfit model had 321 data points(N) and $\chi^2$ = 307.223 (N/$\chi^2$ < 1). The model has 68 free parameters which comprises both biologically relevant model variables, indexes, offsets and scaling factors.

### **S1.5.** Model validation

Using the best fit parameters (supplementary table TS1) we simulated the costimulation experiment where M dose of IFNγ and IL10 are applied simultaneously and the resultant dynamics of STAT1 and STAT3 activation were studied. Due to the inherent parameter uncertainties associated with such biochemical models robust predictions are often not derivable from only one best fit model [11]. So, here we used 40 independently fitted models, each with comparable goodness of fit, to predict the effect of the costimulation on STAT1 and STAT3 dynamics. We achieved quantitative agreements with data for STAT3 and SOCS1 data (Figure 4B and 4D) and STAT1 predictions(Figure S2A) could be quantitatively matched to the data(Figure 4C) by multiplying the predicted STAT1 trajectory to a scaling factor that adjusts the height of the predicted trajectory without further adjustments to the model. Shaded area in Figure 4B, 4C and 4D shows prediction range and the solid lines show the predictions from the fit with lowest $\chi^2$ .

**D. Parameter sampling**

To understand the significance of individual best-fitted parameters on the signaling dynamics of the STATs we sampled the best fit parameters, simulated the model and captured the trajectories of S/1/3.

For a given variable p, varied in the range [lu*p,ub*p] where lb and ub are upper and lower bound of sampling, the following Monte-Carlo sampling method [15] was used

sampled_p = (ub*p - lu*p).* random(1) + lu*p ----- (7)

random(1) generates a random number between 0 to 1. The method was repeated for all the sampled variables and it results in a new parameter vector. We generated 20000 such independent parameter vectors and simulated the model for each parameter vector and analyzed the STATs trajectories. The range of variation allowed for each parameter was a 5 fold in either direction, implying, ub = 5 and lb = 0.2.

**E. Information content calculation**

As implicated in earlier studies[15, 16], we argued, the parameters with maximum information for a desired response type would be the most sensitive parameters and they will be constrained to values in a certain range compared to the less sensitive paramters who in a much wider range can give the response of interest. For instance, if we have 2000 independent parameter sets that results in bell shaped STAT3 responses($Para_{Bell\text{-}shaped}$) to the increasing dose of IFNγ, and 2000 other independent parameter sets resulting in proportional STAT3 dose-responses ($Para_{proportional}$ ), then to calculate the information content of a given parameter m we first calculated the entropy of m in both $Para_{Bell\text{-}shaped}$ and $Para_{proportional}$ .

$$E_{m[Bell\text{-}shaped]} \; ¿ -\sum p(m) \log p(m) \text{---- [8]}$$

where p(m) implies the probability of m to be located in a given bin within the distribution it assumes in the assigned range.

Similarly, $E_{m[proportional]}$ is calculated for the parameter m. The information content of the parameter m is then calculated as[15]

$$I(m) = |E_{m[Bell\text{-}shaped]} - E_{m[proportional]}| \text{ ----- [9]}$$

Information content of all the model variables were calculated in a similar manner. This method was also used to calculate the information content and identify the most sensitive parameter(s) determining the robustness of IL10-STAT3 signaling during costimulation.
