## Supplementary figures and images for "Regulation of distinct STAT3 dynamics in pro (IFNγ) and anti (IL-10) inflammatory pathways and in their cross-talk: insights from a data-driven model"

### Supplementary figures S1 to S3

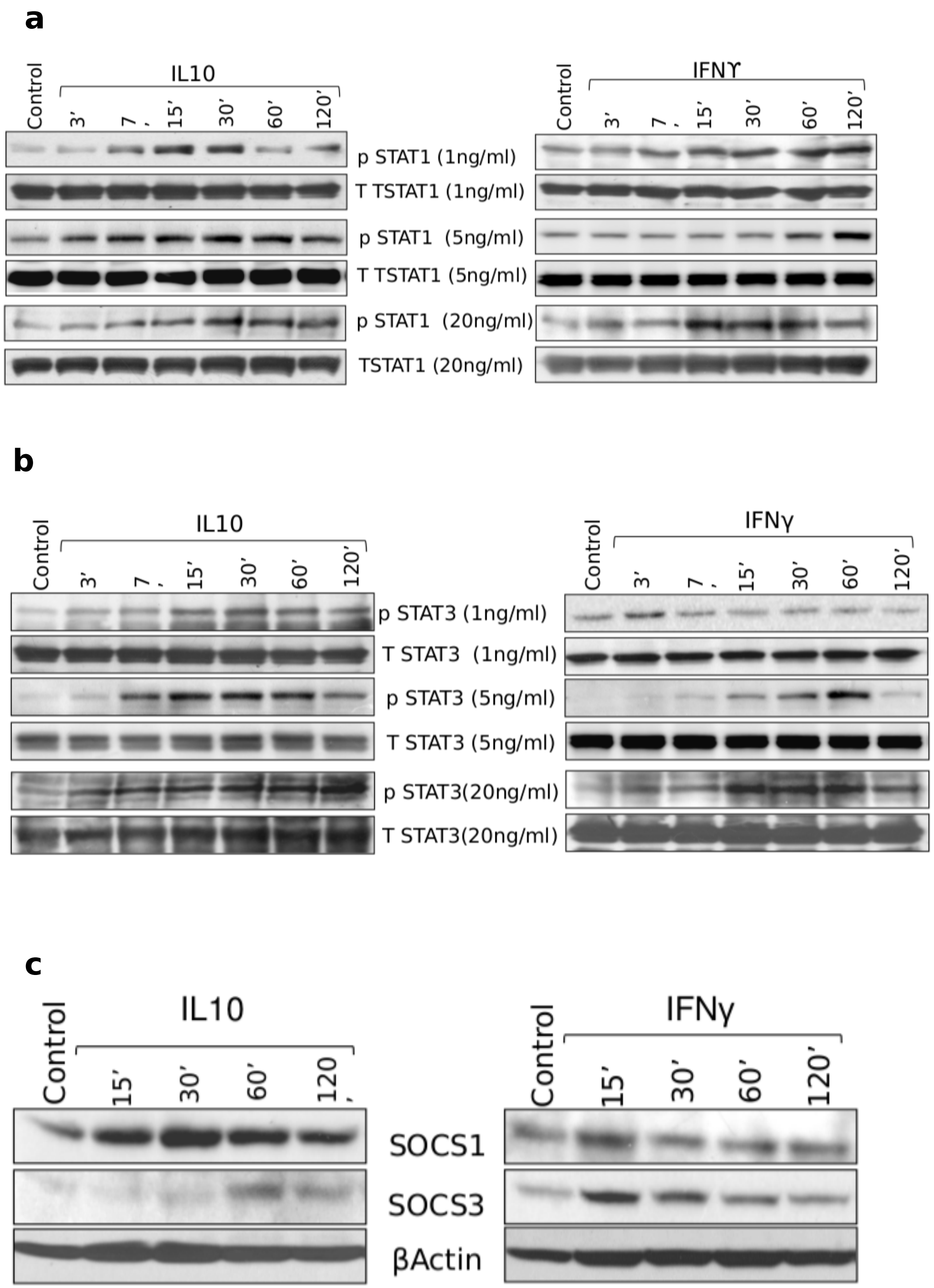

a

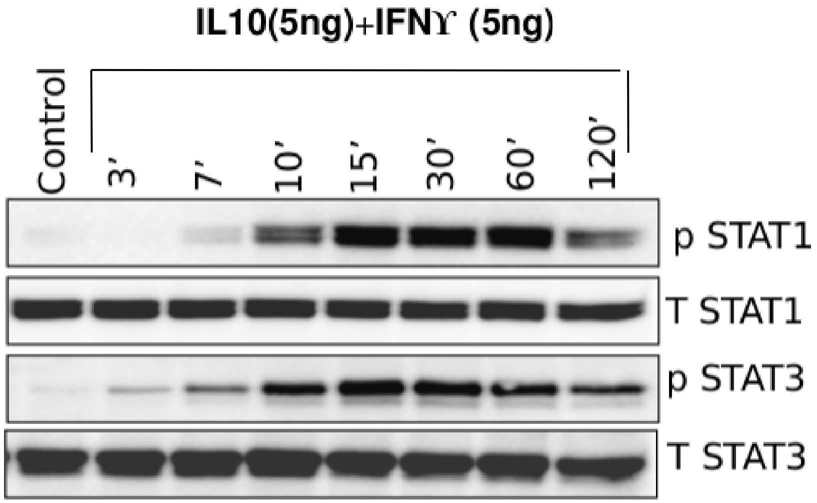


b

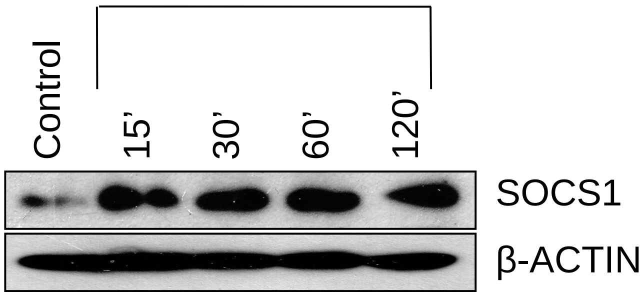


c

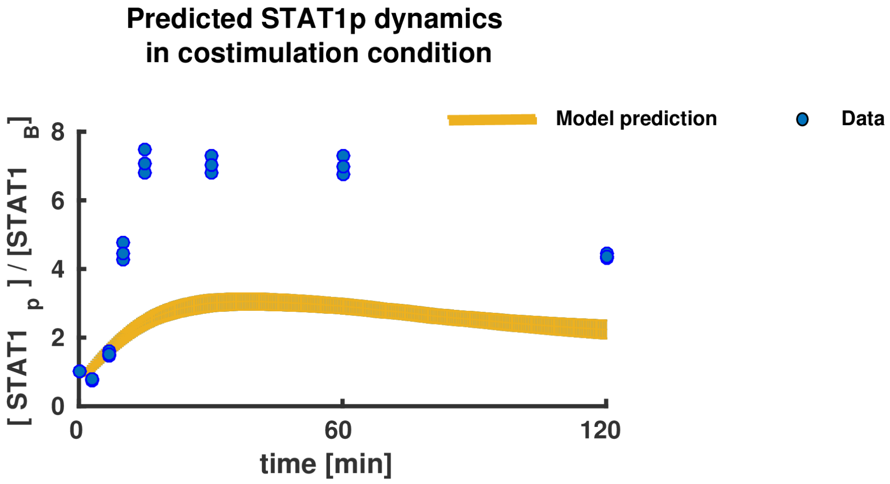


Figure S2

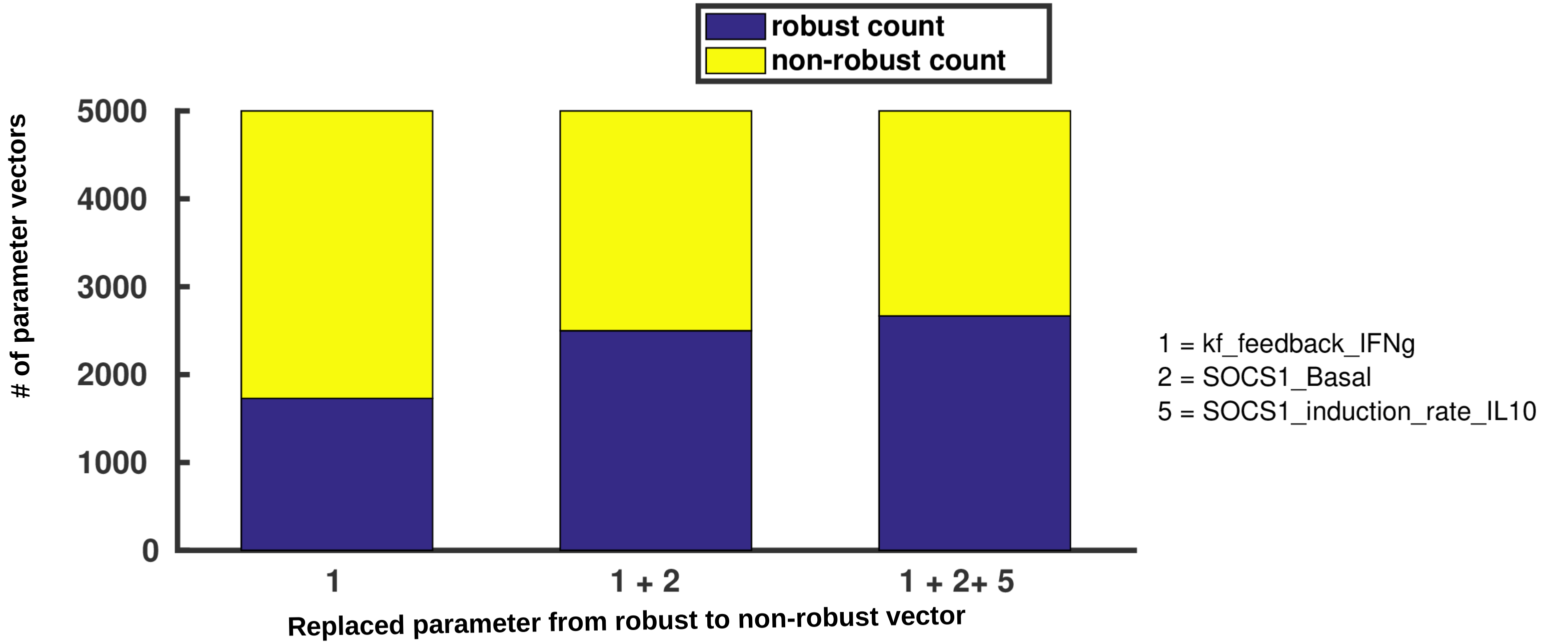
